# Conditional Sequence-Structure Integration: A Novel Approach for Precision Antibody Engineering and Affinity Optimization

**DOI:** 10.1101/2024.07.16.603820

**Authors:** Benyamin Jamialahmadi, Mahmood Chamankhah, Mohammad Kohandel, Ali Ghodsi

## Abstract

Antibodies, or immunoglobulins, are integral to the immune response, playing a crucial role in recognizing and neutralizing external threats such as pathogens. However, the design of these molecules is complex due to the limited availability of paired structural antibody-antigen data and the intricacies of structurally non-deterministic regions. In this paper, we introduce a novel approach to designing antibodies by integrating structural and sequence information of antigens. Our approach employs a protein structural encoder to capture both sequence and conformational details of antigen. The encoded antigen information is then fed into an antibody language model (aLM) to generate antibody sequences. By adding cross-attention layers, aLM effectively incorporates the antigen information from the encoder. For optimal model training, we utilized the Causal Masked Language Modeling (CMLM) objective. Unlike other methods that require additional contextual information, such as epitope residues or a docked antibody framework, our model excels at predicting the antibody sequence without the need for any supplementary data. Our enhanced methodology demonstrates superior performance when compared to existing models in the RAbD benchmark for antibody design and SKEPMI for antibody optimization.

## 1 Introduction

In the complex and crucial domain of protein design, antibody design stands out as a particularly challenging task due to the sophisticated structures and specific antigen-binding capabilities of antibodies. Antigens, molecules that trigger an immune response, are typically found on the surfaces of pathogens such as bacteria, viruses, and fungi and are specific to each pathogen. These antigens are crucial targets for antibodies, or immunoglobulins, distinct proteins produced by the immune system to recognize and neutralize these pathogens. As the primary defense mechanism of the body, antibodies play a pivotal role in the adaptive immune response, ensuring specificity and memory against pathogens encountered previously (Janeway et al., 2001).

Antibodies are structurally similar to a Y-shape, comprising two identical heavy chains and two identical light chains, which together form a functional unit poised for antigen binding (Figure 1). The antigen-binding site is largely defined by the Complementarity-Determining Regions (CDRs) within these chains (Schroeder and Cavacini, 2010). The third CDR of the heavy chain (H3) exhibits remarkable variability, allowing antibodies to attach to many different types of harmful substances, thus determining the specificity of the immune response (Al-Lazikani et al., 1997).

**Figure 1:**
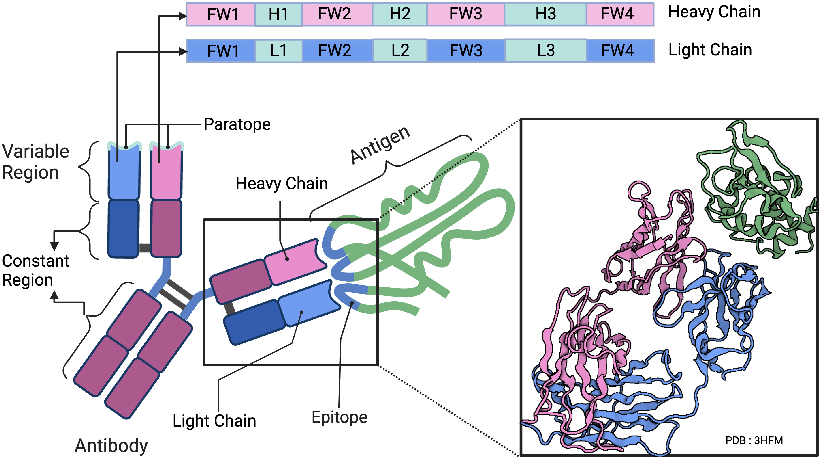
Schematic and structural representation of the antibody’s typical Y-shaped formation, consisting of two identical heavy chains and two identical light chains. Each chain is composed of alternating framework regions (FW) and complementarity-determining regions (CDRs). There are three CDRs (H1, H2, H3 for the heavy chain and L1, L2, L3 for the light chain) that are instrumental in binding to specific parts of the antigen, and four framework regions (FW1, FW2, FW3, FW4) that support the overall structure. The paratope, formed by the CDRs, engages the antigen’s epitope in a precise lock-and-key manner. The epitope is a specific part of an antigen that is recognized and bound by an immune system component. The right side of the figure showcases the three-dimensional configuration of an antibody-antigen complex (PDB: 3HFM), highlighting the interaction between the antibody (in pink and blue) and the antigen (in green), with a focus on how the paratope’s CDRs closely contact with the antigen’s epitope. Adapted from “Antigen Recognition by Antibodies”, by BioRender.com (2023). Retrieved from https://app.biorender.com/biorender-templates.

The central task of antibody design, a crucial aspect of therapeutic development, involves the prediction of the variable region sequences of an antibody, which directly interact with antigens. This prediction is based on the detailed structure and sequence of a specific antigen. Achieving this level of specificity in antibody design is a complex task due to the immense diversity of possible antibody sequences and the intricate three-dimensional structures they can form. Consequently, traditional approaches involving complex physics-based calculations have become less efficient (Li et al., 2014a; Adolf-Bryfogle et al., 2018).

Advancements in computational techniques have opened new avenues in antibody design, particularly through the adoption of deep learning methods. These methods focus on enhancing the design accuracy by integrating both the sequence and the structural aspects of antibodies (Jin et al., 2021; Luo et al., 2022a; Kong et al., 2022a; 2023). However, these innovative models encounter substantial challenges, which often slow down the creation of accurate and widely applicable models. The key challenges are described below:

### Limited availability of paired structural data

The main obstacle in antibody design is the shortage of extensive antibody-antigen structural data for training deep learning models. For instance, the Structural Antibody Database (SAbDab) dataset Dunbar et al. (2014), a widely-used and comprehensive resource containing paired antibody-antigen structures, includes only about 5,000 samples. This lack of data limits the models’ overall efficacy and applicability by making it more difficult for the deep learning models to learn from and generalize across a wide variety of antigens.

### Impact of CDR flexibility on sequence prediction

The structural flexibility of CDRs (Jeliazkov et al., 2018) presents a significant challenge in accurately predicting their sequences. Zheng et al. (2023) emphasize that in such flexible regions, there is a weaker correlation between residue identities and their structural context. This aspect is particularly problematic in methods attempting to design both the sequence and structure of antibodies simultaneously, often leading to errors and resulting in the generation of inaccurate sequences.

### Dependency on contextual information

Existing models in antibody design often rely on additional data, such as the structural arrangements of antibodies within their target environment (docked antibody frameworks) Luo et al. (2022b) or the specific shapes and conformations of antigenic epitopes Kong et al. (2023). Although this contextual information plays a vital role in enhancing the scalability and overall effectiveness of the aforementioned methods, it is often challenging to obtain.

To address the above challenges, we propose Aligned Integrated Design for Antibodies (AIDA), a novel approach that leverages the advancement of protein language models, which have shown promise in forecasting protein structures and aiding in protein design. At its core, AIDA is driven by an antibody language model (aLM) and employs a specialized protein encoder to capture the sequential and structural information of antigens. Importantly, it integrates antigen adapter modules into the aLM, facilitating efficient information transfer from the encoded antigen to the aLM, which is crucial for the conditional prediction of antibody sequences.

AIDA’s emphasis on sequence prediction skillfully circumvents the structurally non-deterministic nature of antibodies. It capitalizes on pre-trained models, which have undergone extensive training on large unsupervised datasets, effectively addressing the data scarcity challenge in antibody design. The efficacy of AIDA was validated in the Rosetta Antibody Design (RAbD) benchmark (Adolf-Bryfogle et al., 2018), where it demonstrated superior performance in recovering antibody sequences, surpassing existing state-of-the-art models without the reliance on additional contextual information.

### 1.1 Related Works

#### Computational antibody design

is an expanding field that uses various methods to predict antibody sequences for a given antigen. Traditional approaches often involve optimizing complex energy functions. Additionally, these approaches require accurate simulation of real-world protein interactions, which is extremely complicated (Pantazes and Maranas, 2010; Li et al., 2014b; Lapidoth et al., 2015; Adolf-Bryfogle et al., 2018; Warszawski et al., 2019). This complexity is due to the fact that the full nature of protein-protein interactions is too intricate to be captured entirely by statistical functions Graves et al. (2020). In response, deep learning has become increasingly prominent, branching into two main approaches: sequence-based and structure-sequence co-design methodologies.

Sequence-based models, such as the ones proposed by Liu et al. (2019); Saka et al. (2021); Akbar et al. (2022); Melnyk et al. (2023), focus on using deep learning to develop antibodies by modeling the one-dimensional sequence data. However, these methods can underperform as they disregard structural information. On the other hand, co-design methods, as seen in the works of Jin et al. (2021); Kong et al. (2022b; 2023); Luo et al. (2022c); Verma et al. (2023), address antibody design by using graph neural networks or diffusion models to develop antibody sequences and 3D structures simultaneously. Despite their use of structural data, these methods still face the challenges mentioned in §1.

Our model addresses these limitations by taking a middle ground between these approaches. We employ graph modeling to encode both the sequence and structural information of the antigen. Then, we utilize sequence modeling to decode this information into the corresponding antibody sequence, effectively balancing the need for structural data with the advantages of sequence prediction.

#### Protein Structural Encoding

The advancement of structure-based methodologies in computational biology has been significantly driven by innovations in encoding spatial characteristics of protein structures. Initially, these methodologies utilized 3D Convolutional Neural Networks (CNNs), as introduced by Derevyanko et al. (2018), and later incorporated Graph Neural Networks (GNNs), evidenced by the work of Gligorijević et al. (2021), Baldassarre et al. (2021), Jing et al. (2021), Wang et al. (2023), and Aykent and Xia (2022). This field has recently seen a surge in exploring the potential of pre-training structural encoders on extensive and unlabeled datasets. Pioneering contributions in this area, such as those by Hermosilla and Ropinski (2022); Chen et al. (2023); Guo et al. (2022), have adopted approaches such as contrastive learning, self-prediction, and denoising score matching. Additionally, Zhang et al. (2023) have proposed the Geometry-Aware Relational Graph Neural Network (GearNet), utilizing relational graph convolutional layers and edge message passing with augmentation functions, offering a novel perspective in self-supervised learning for protein structures. These developments are expanding the boundaries of what can be achieved in the encoding of protein structural information.

#### Protein/Antibody Language Models

Protein language models (pLMs) such as ESM Lin et al. (2022) and ProtTrans Elnaggar et al. (2021), which have been pre-trained using the Masked Language Model (MLM) technique, have significantly enhanced our understanding of protein sequences. These advancements have motivated researchers to use similar methods in analyzing antibody sequences, leading to the creation of specialized models such as AbLang Tobias H. Olsen and Deane (2022), AntiBERTa Leem et al. (2022), AntiBERTy Ruffolo et al. (2021), and BALM Jing et al. (2023). AbLang, trained on the OAS database Olsen et al. (2022), focuses on filling in missing residues in B-cell receptor repertoire sequences. AntiBERTa, utilizing a 12-layer transformer architecture (Vaswani et al., 2023), provides a nuanced numeric representation of antibody sequences, capturing essential elements of antibody functionality, and is adaptable for paratope prediction tasks. AntiBERTy, on the other hand, groups antibodies into clusters that mimic the process of affinity maturation. Meanwhile, BALM stands out for its ability to predict both the function and structure of antibodies, exemplifying the significant role of machine learning in the field of immunology.

## 2 Method

In this section, we first outline the task formulation, followed by a detailed exposition of our proposed method, ensuring a logical flow from problem understanding to solution presentation.

### 2.1 Task Formulation

The objective is to predict antibody sequence *b* consisting of light and heavy chains *l* and *h*, respectively, given the input antigen information *g*. In this formulation, *g* is represented by a sequence-structure tuple (*s*_*i*_, x_*i*_) for *i* = 1, …, *n*, where *s*_*i*_ ∈ 𝒜, the set of all naturally occurring amino acids, and x_*i*_ ∈ ℝ^3^ is the coordinates of the alpha carbon atom of amino acid at position *i*. The task aims to learn parameters *θ* to maximize the conditional probability:

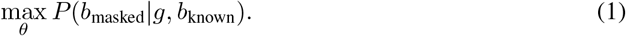

In specific cases, part of the antibody sequence, typically the framework, might already be identified, indicated as *b*_known_. Our goal is to accurately predict the unspecified segments of the antibody sequence, referred to as *b*_masked_. For the case of full antibody prediction, no section is known, hence *b*_known_ = ∅.

### 2.2 Architecture

Informed by our previous discussions, we elected to concentrate solely on the design of the antibody sequence. Our objective is to gain a deeper understanding of the antigen to create a complementary antibody sequence effectively. To achieve this, we adopted an encoder-decoder architecture.

The encoder in our framework is tasked with capturing the antigen’s structural and sequential details comprehensively. The decoder, on the other hand, uses this information to produce the corresponding antibody sequence, systematically incorporating all essential aspects of the antigen. We utilized GearNet, a structural encoder pre-trained on the AlphaFold dataset (Figure 2A), for antigen encoding. This pre-training process equipped GearNet with a robust understanding of protein structures, making it adept at accurately encoding the information of antigens. The encoder function ENC(*g*) produces *H*_*g*_, which is in the shape of *n* × *d*_GearNet_, where *n* is the length of the antigen sequence and *d*_GearNet_ is the dimensionality of the encoded space. *H*_*g*_ is then passed through an adapter module, undergoing a linear transformation with weights *W*_AdaptEnc_ to produce *E*_*g*_ such that *E*_*g*_ = *W*_AdaptEnc_ · *H*_*g*_, and we parameterize the dimensionality of *E*_*g*_ with *d*_AdaptEnc_ = 256. The transformed representation *E*_*g*_ is then passed to the decoder, serving as the foundation for generating corresponding antibody sequences and translating the complex antigen information into a usable format for antibody design.

**Figure 2:**
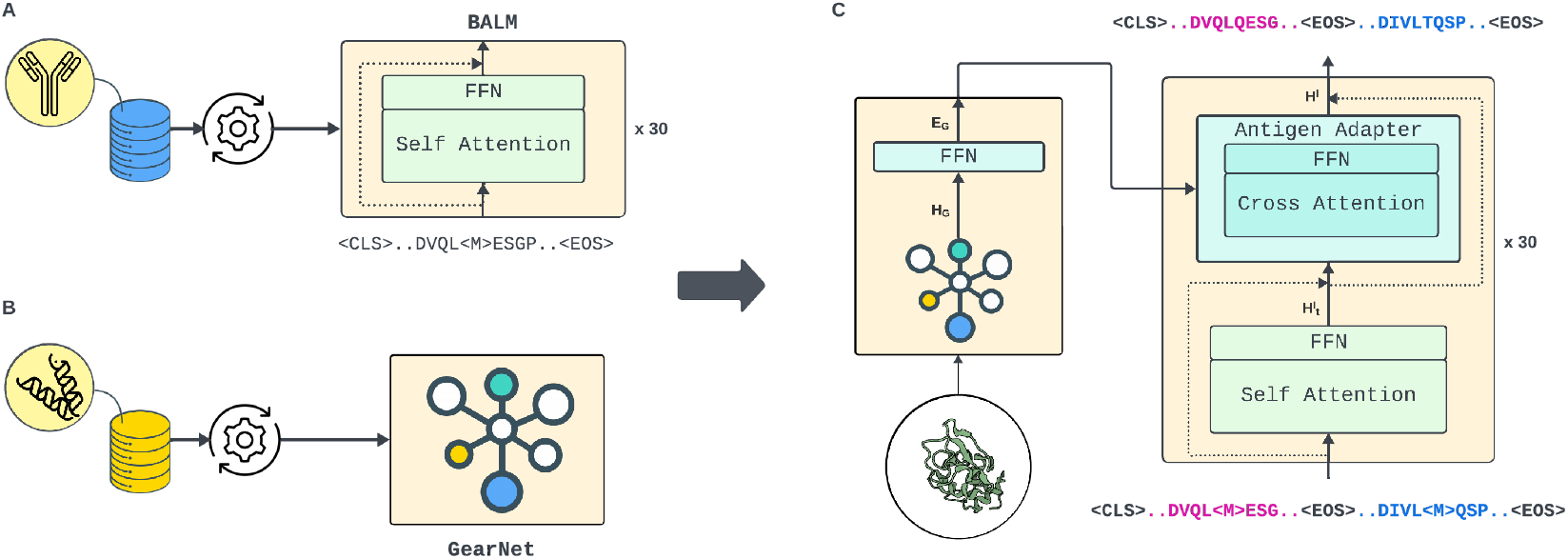
This diagram showcases our proposed encoder-decoder model for antibody sequence prediction, combining the strengths of two pre-trained networks. Panel A features BALM, a transformer-based model pre-trained on a vast dataset of single-chain antibodies using the MLM objective for sequence processing. In Panel B, GearNet is presented as a general-purpose protein structure encoder, pre-trained on the comprehensive AlphaFold dataset to encode structural data into graph form. Panel C unifies these components: GearNet’s graph-encoded antigen information is channeled into BALM via an antigen adapter, using cross-attention mechanisms to translate intricate antigen structures into corresponding antibody sequences.

Our decoding component, the Bio-inspired Antibody Language Model (BALM), leverages a transformer-based self-attention mechanism with 30 layers and rotary positional encoding to understand the unique and conserved properties of antibodies (Figure 2B). Initially trained on a large dataset of unlabeled antibody sequences, BALM effectively captures contextual embeddings essential for inferring binding functions. Notably, BALM employs a unique antibody positional encoding method based on the IMGT numbering scheme (Lefranc et al., 2003), which provides a consistent framework for identifying amino acid positions, thereby enhancing its ability to generate meaningful embeddings.

In our methodology, the antibody sequence representation provided to BALM adopts a structure similar to ESM-2; it commences with a [CLS] token followed by the antibody’s heavy chain sequence, a [EOS] token, the antibody’s light chain sequence, and another [EOS] token. To adapt BALM for conditional generation crucial to our approach, we integrated antigen adapter layers after each self-attention layer within BALM (Figure 2C). This integration facilitates the efficient incorporation of the encoded antigen information into the decoding process. Each antigen adapter layer consists of a cross-attention layer followed by a feed-forward layer. In the cross-attention operation denoted as CrossAtt(Q, K, V), the output of each block in adapted BALM is given by:

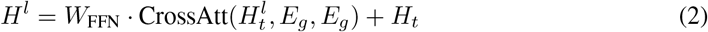

where *l* signifies the layer number, *E*_*g*_ represents the encoded antigen information, and *W*_FFN_ are the weights of the feed-forward layer, and *H*_*t*_ is the output from the previous feed-forward and self-attention layers. This refined structure ensures a synergistic interplay between the antigen’s structural and sequential information, aligning with our objective of generating corresponding antibody sequences based on the encoded antigen information.

### 2.3 Training and Inference

During the training phase, our model utilizes the Causal Masked Language Modeling (CMLM) objective (Ghazvininejad et al., 2019), which facilitates the non-autoregressive generation of antibody sequences informed by antigen structure. A subset of tokens from the antibody sequence *B* is replaced with a [MASK] token, with the number of masked tokens uniformly sampled between 1 and |*B*|, where |*B*| denotes the sequence length. To direct the model’s learning toward the variable and critical CDRs rather than the more conserved framework regions, we apply Focal Loss Lin et al. (2018) as our training loss function. Focal Loss is mathematically formulated as:

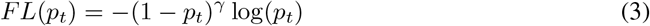

where *p*_*t*_ is the model’s estimated probability for the class label *t* and *γ* (In our training, *γ* = 2) is the focusing parameter that adjusts the rate at which easy-to-classify residues are down-weighted. This loss function is particularly effective in managing class imbalance and helps the model focus on the challenging CDR regions essential for our task.

During inference, we strategically place the [MASK] token at positions corresponding to the parts of the sequence we aim to predict, such as the H3 region. These masked positions are then predicted using our adapted BALM model. Unlike the iterative mask prediction strategy suggested in Ghazvininejad et al. (2019), which incrementally refines the sequences, our observations indicated no significant improvements with multiple iterations. Consequently, we opted for a single iteration of unmasking, which efficiently yields accurate sequence predictions without the need for iterative refinement.

The training of our model was conducted using the Adam optimizer on one NVIDIA A40 GPU, paired with an inverse square root learning rate scheduler. This scheduler featured a warm-up phase of 1,000 steps, after which the learning rate peaked at 10^*−*4^. To preserve the integrity of the pre-trained models, we initially kept all pre-trained weights frozen during the first ten epochs. After this period, we unfroze the pre-trained weights and set their learning rate to a tenth of that of the newly added layers. Importantly, we initialized *W*_FFN_ in antigen adapters to zeroes. This weight initialization ensures that the newly added layers do not immediately disrupt the predictive capabilities of BALM, allowing the model to maintain its pre-trained performance while the new layers gradually adapt during the fine-tuning process. This approach allowed for end-to-end training, providing the model with the opportunity to adjust to the specific characteristics of our dataset gradually.

## 3 Experiment

In this section, we rigorously evaluate our proposed model through four experiments, benchmarking AIDA against established models in antibody design. The experiments include (1) Single CDR design (§3.1), (2) Multiple CDR design (§3.2), (3) Full antibody design (§3.3), and (4) Antibody Optimization (§3.4).

To thoroughly evaluate our model, we compared it against leading models in both sequence-based and co-design categories. In the sequence-based category, we included **ReprogBert** (Melnyk et al., 2023), which reprograms a pre-trained English BERT model for antibody design. Notably, ReprogBert utilizes IgFold (Ruffolo et al., 2023) for generating antibody structures and calculating Root Mean Square Deviation (RMSD). In the co-design category, our comparisons included **DiffAb** (Luo et al., 2022b), employing a diffusion process for simultaneous sequence and structure prediction, **AbODE** (Verma et al., 2023), modeling antigen-antibody interactions via graph Partial Differential Equations (PDE), and **dyMEAN** Kong et al. (2023), which features an adaptive multi-channel encoder for full atom modeling and a ’Shadow Paratope’ technique for comprehensive antigen interaction analysis.

To better equip BALM for the specific requirements of our experiments and to enhance its understanding of the interplay between heavy and light chains in antibodies, we drew upon the strategies of Burbach and Briney (2023). Their work significantly boosted the performance of their aLM by focusing on training with natively paired heavy and light chain sequences. Motivated by this, we initiated a secondary pre-training phase for BALM, leveraging the OAS database to curate paired antibody sequences. These sequences were clustered based on a 50% sequence identity threshold, yielding a collection of approximately 100,000 non-redundant antibody sequences. For this pre-training stage, we implemented an MLM objective supplemented by entropy-guided masking, as recommended by Jing et al. (2023). This pre-training phase is aimed at substantially enriching BALM’s comprehension of antibody sequences and the complex relationship between heavy and light chains.

### 3.1 Single CDR Design

In this experiment, we tackled the challenge of predicting a single CDR, given the other domains of the light and heavy chains. We used a masking-unmasking technique, where the target CDR is initially masked, and the model then predicts its sequence. This process tests the model’s ability to integrate contextual information from the known sections of the antibody’s variable region to design the target CDR.

To train our model, we utilized the SAbDab dataset, which includes about 5,000 samples of paired antibodies and nanobodies. We clustered samples based on their CDR-H3 sequences using the MMseqs2 algorithm and the BLOSUM 62 scoring matrix, at a 40% sequence identity threshold. Sixty samples from the RAbD benchmark were reserved for testing, and clusters containing these test samples were excluded from the training and validation sets to prevent data leakage. From the remaining clusters, 10% were randomly selected for validation, with the rest used for training.

The model’s predictive performance was assessed using three main metrics: Amino Acid Recovery (**AAR**), Contact Amino Acid Recovery (**CAAR**), and Root Mean Square Deviation (**RMSD**). AAR measures the accuracy of the predicted amino acid sequence, while CAAR evaluates the model’s ability to predict residues that interact with the antigen. As our model does not directly predict protein structure, we utilized the dyMEAN structure prediction model, available on its official GitHub repository, to predict the complex tertiary structure for the generated antibody sequences. We then used RMSD to measure structural accuracy, calculating the average deviation between the backbone atoms of the predicted and actual protein structures post-alignment using the Kabsch algorithm. A lower RMSD indicates higher structural fidelity. By combining AAR and CAAR for sequence accuracy and RMSD for structural accuracy through dyMEAN predictions, we comprehensively evaluate our model’s efficacy in replicating both the antibody sequences and their functional conformations.

#### Results

As indicated in Table 1, our AIDA model demonstrated remarkable proficiency in predicting the sequence of a single CDR. This success highlights AIDA’s ability to use context from conserved antibody regions and integrate targeted antigen information through adapter modules, enabling precise engineering of CDR regions critical for antigen recognition. Notably, AIDA outperformed models like DiffAb, AbODE, and dyMEAN in designing the heavy chain’s third CDR (H3), showcasing its advanced sequence prediction capabilities. While AIDA excelled in both sequence-based and most structure-based metrics, it is important to consider potential inaccuracies from the external structure prediction tool used for evaluation.

**Table 1:** Comparison of AIDA with existing models (ReprogBert, Diffab, AbODE, dyMEAN) in single CDR design. Metrics: Amino Acid Recovery (AAR), Contact Amino Acid Recovery (CAAR), and Root Mean Square Deviation (RMSD). AIDA shows superior performance in H1, H2, H3 AAR, and CAAR.

| Model | H1 |  |  | H2 |  |  | H3 |  |  |
| --- | --- | --- | --- | --- | --- | --- | --- | --- | --- |
|  | AAR↑ | CAAR↑ | RMSD↓ | AAR↑ | CAAR↑ | RMSD↓ | AAR↑ | CAAR↑ | RMSD↓ |
| ReprogBert | - | - | - | - | - | - | 36.30% | - | 2.47 |
| Diffab | 68.00% | 60.02% | 0.55 | 54.64% | 42.77% | **0.41** | 37.47% | 20.88% | 2.08 |
| AbODE | 70.50% | - | 0.65 | 55.70% | - | 0.73 | 39.80% | - | 1.73 |
| dyMEAN | 76.55% | 63.19% | 0.56 | 69.52% | 63.50% | 0.48 | 43.20% | 27.83% | **1.58** |
| **AIDA** | **80.56%** | **69.44%** | **0.47** | **71.25%** | **66.46%** | 0.48 | **46.41%** | **33.04%** | 1.68 |
| Model | L1 |  |  | L2 |  |  | L3 |  |  |
|  | AAR↑ | CAAR↑ | RMSD↓ | AAR↑ | CAAR↑ | RMSD↓ | AAR↑ | CAAR↑ | RMSD↓ |
| Diffab | 45.88% | 39.24% | 0.55 | 54.68% | 25.00% | **0.11** | 43.99% | 25.32% | **0.71** |
| dyMEAN | **74.11%** | **65.49%** | **0.48** | **81.98%** | **51.94%** | 0.20 | **51.93%** | 33.33% | 0.87 |
| **AIDA** | 67.01% | 58.66% | 0.51 | 78.09% | 49.17% | 0.65 | 47.03% | **62.07%** | 1.63 |

### 3.2 Multiple CDR Design

In this experiment, we predicted the sequences of all six CDRs (three from the heavy chain and three from the light chain) simultaneously, given the antigen structure and the antibody framework sequence. This demonstrated AIDA’s robustness in handling complex design tasks, integrating and processing intricate antigen information, and managing multiple CDRs. The evaluation followed the same setting as the single CDR design experiment (§3.1).

#### Results

The results in Table 2 show AIDA’s exceptional capability in handling complex antibody design tasks. AIDA maintained consistent performance in multiple CDR design, with less dependency on known antibody sections compared to other methods like dyMEAN, which saw a significant drop in performance, especially in H3 design. AIDA’s higher AAR and CAAR, along with significantly lower RMSD, underscore its superiority in accurate antibody sequence prediction.

**Table 2:** Comparison of AIDA with existing models in multiple CDR design. Metrics: AAR, CAAR, RMSD. AIDA excels in AAR and CAAR for L3 and heavy chain CDRs, outperforming Diffab and dyMEAN overall.

| Model | AAR↑ |  |  |  |  |  |  |  |  |
| --- | --- | --- | --- | --- | --- | --- | --- | --- | --- |
|  | L1 | L2 | L3 | H1 | H2 | H3 |  |  |  |
| Diffab | 52.02% | 56.03% | 47.88% | 68.97% | 54.64% | 39.82% |  |  |  |
| dyMEAN | **75.89%** | **83.65%** | 52.66% | 76.13% | 68.48% | 37.51% |  |  |  |
| **AIDA** | 63.20% | 75.4% | **61.49%** | **79.76%** | **70.23%** | **44.98%** |  |  |  |
| Model | CAAR↑ |  |  |  |  |  | AAR↑ | Overall CAAR↑ | RMSD↓ |
|  | L1 | L2 | L3 | H1 | H2 | H3 |  |  |  |
| Diffab | 46.73% | 28.06% | 27.22% | 61.85% | 44.01% | 24.31% | 51.26% | 40.35% | 1.95 |
| dyMEAN | **67.99%** | **53.61%** | 35.26% | 63.11% | 62.52% | 23.11% | 60.36% | 50.30% | 1.35 |
| **AIDA** | 58.36% | 44.72% | **46.30%** | **67.50%** | **65.74%** | **31.89%** | **63%** | **53.67%** | **1.01** |

### 3.3 Full Antibody Prediction (Whole Variable Region Prediction)

In this critical phase of our research, we extended the scope of AIDA’s capabilities by challenging it to predict the entire variable region of antibodies. This includes both the heavy and light chains, encompassing all the CDRs and framework regions. Unlike the previous tasks that focused on specific CDRs, this task requires the model to generate a more extensive and complete sequence, reflecting the full complexity of an antibody’s variable region. This experiment included similar settings as the single CDR (§3.1).

#### Results

Table 3 shows the results of the full antibody prediction experiment. Our model was compared against dyMEAN, the only other method capable of addressing this extensive task. Remarkably, AIDA surpassed dyMEAN in all sequential metrics for predicting the entire variable region of the antibody. This indicates AIDA’s superior ability to model the interaction of antigens and antibodies.

**Table 3:** Performance of AIDA in predicting the complete variable region of antibodies, including heavy and light chains and all CDRs, compared against dyMEAN. Metrics such as Amino Acid Recovery (AAR), Contact Amino Acid Recovery (CAAR), and Root Mean Square Deviation (RMSD) highlight AIDA’s superior predictive capabilities in this extensive and complex task.

| Model | AAR | CAAR | RMSD |
| --- | --- | --- | --- |
| dyMEAN | 70.35% | 40.02% | **1.11** |
| AIDA | **72.48%** | **45.15%** | 1.21 |

### 3.4 Antibody Optimization

In our antibody optimization experiment, we aimed to enhance the binding affinity of antibodies to specific antigens, focusing on the H3 region. The models were trained on the SAbDab dataset, and SKEMPI V2.0 Jankauskaitė et al. (2018) was used for testing under the same settings as the single CDR design experiment (§3.1). Performance was assessed using two metrics: (1) the binding affinity change (ΔΔ*G*), determined through a GNN-based predictor Shan et al. (2022) as recommended by Kong et al. (2023), and (2) the average number of modified residues (Δ*L*), indicating the importance of minimal sequence alterations for improved affinity Ren et al. (2022).

To tailor our model for this experiment and control the number of modified residues, we masked *n* random tokens from the H3 region for AIDA to predict. We then used the dyMEAN structure prediction model to predict the antibody-antigen complex structure. Both models generated 100 sequences for each sample, selecting the one with the best ΔΔ*G* for metric calculation.

#### Results

In Table 4, the results demonstrate AIDA’s effectiveness in enhancing antibody binding affinity compared to dyMEAN. The evaluation, using ΔΔ*G* and Δ*L*, shows that AIDA, particularly in its unrestricted form, significantly outperforms dyMEAN. AIDA achieves a notable improvement in binding affinity with ΔΔ*G* = −5.48 while maintaining a reasonable degree of residue modification (Δ*L* = 4.10), highlighting its capability to optimize antibody sequences for higher affinity with minimal alterations.

**Table 4:** Comparison of AIDA and dyMEAN in Antibody H3 Region Optimization: This table illustrates the performance of AIDA and dyMEAN in enhancing the binding affinity of the H3 region of antibodies. Metrics used for evaluation include the change in binding affinity (ΔΔ*G*) and the average number of modified residues (Δ*L*). The dyMEAN-*n* and AIDA-*n* variants indicate models restricted to modifying only *n* tokens. In contrast, dyMEAN and AIDA represent configurations where *n* tokens, ranging from 1 to the length of the H3 region, were randomly altered. For each sample, both models generated 100 sequences, with the best-performing sequence chosen for evaluation. These results highlight the models’ effectiveness in achieving enhanced binding affinity with minimal sequence modifications.

| Method | $\Delta\Delta G \downarrow$ | $\Delta L \uparrow$ |
| --- | --- | --- |
| dyMEAN-1 | -2.97 | 1.00 |
| dyMEAN-2 | -3.22 | 1.35 |
| dyMEAN-4 | -3.40 | 2.22 |
| dyMEAN-8 | -3.84 | 4.84 |
| dyMEAN | -3.27 | 4.20 |
| AIDA-1 | -3.47 | 0.82 |
| AIDA-2 | -4.46 | 1.47 |
| AIDA-4 | -5.24 | 2.27 |
| AIDA-8 | -3.48 | 4.12 |
| AIDA | **-5.48** | 4.10 |

### 3.5 Ablation Study

In our ablation study, we assess the necessity and value of the following components in the AIDA model: Focal Loss, the second paired pre-training process for BALM, CMLM training (replaced with a full masking strategy for training), and the antigen adapter (where we only fine-tuned BALM). The results (Table 5) across the H3, All CDRs, and Full Antibody design settings show that each component plays a crucial role in the model’s performance. AIDA achieves the highest AAR and CAAR scores in H3 and Full Antibody settings, demonstrating the effectiveness of these components, particularly in challenging regions. Interestingly, CMLM performs best in All CDRs, suggesting advantages in general binding site prediction. Overall, the combination of these components is crucial for maximizing AIDA’s accuracy and consistency across antibody regions.

**Table 5:** Ablations of different components in AIDA.

| Model | H3 |  | All CDRs |  | Full antibody |  |
| --- | --- | --- | --- | --- | --- | --- |
|  | AAR↑ | CAAR↑ | AAR↑ | CAAR↑ | AAR↑ | CAAR↑ |
| AIDA | **46.41%** | **33.04%** | 63.00% | 53.67% | **72.48%** | **45.15%** |
| Focal | 46.01% | 31.86% | 61.94% | 52.62% | 72.10% | 43.26% |
| Paired Pretraining | 45.32% | 31.84% | 63.23% | 53.82% | 71.58% | 43.49% |
| CMLM | 44.12% | 30.37% | **64.14%** | **54.71%** | 71.91% | 43.54% |
| Antigen Adapter | 43.10% | 29.87% | 61.15% | 57.16% | 72.07% | 42.20% |

## 4 Limitations and Future Work

In advancing computational antibody design with AIDA, we’ve identified key limitations. One major challenge in this field is the lack of computational metrics that accurately reflect antibody-antigen interactions, suggesting a significant area for future research.

Additionally, the scarcity of antigen-antibody paired data limits our model’s predictive accuracy despite using pre-trained models and transfer learning. Future work could explore increasing sample numbers through wet lab experiments or data augmentation.

Finally, AIDA’s precision is dependent on the structural models used for antigen encoding; inaccuracies here can affect the final design. However, our model’s modularity allows for future improvements by incorporating advanced protein structure encoders focused on surface analysis and binding sites.

## 5 Conclusion

In conclusion, our study contributes significantly to the field of computational antibody design, tackling the prevalent challenges of data scarcity and structural flexibility in antibody regions. By introducing Aligned Integrated Design for Antibodies (AIDA), which utilizes an encoder-decoder architecture integrating both structural and sequence information of antigens, we have made noteworthy progress in antibody sequence prediction. Our proposed model exhibited enhanced performance over established methods such as dyMEAN and Diffab in different experiments, including predicting the single and multiple CDRs, the entire variable region of antibodies, and optimizing the antibodies’ binding affinity. AIDA’s focus on sequence prediction and its effective management of complex antigen information marks a meaningful step forward in the development of efficient and accurate methods for antibody design, opening new avenues for advancements in biomedical research and healthcare applications.

## NeurIPS Paper Checklist

### 1. Claims

Question: Do the main claims made in the abstract and introduction accurately reflect the paper’s contributions and scope?

Answer: [Yes]

Justification: The abstract and introduction clearly state the main claims of the paper, including the introduction of a novel approach (AIDA) for antibody design that integrates structural and sequence information of antigens. The claims made are supported by theoretical and experimental results presented in the paper, demonstrating AIDA’s superior performance compared to existing models in various benchmarks.

Guidelines:

- The answer NA means that the abstract and introduction do not include the claims made in the paper.
- The abstract and/or introduction should clearly state the claims made, including the contributions made in the paper and important assumptions and limitations. A No or NA answer to this question will not be perceived well by the reviewers.
- The claims made should match theoretical and experimental results, and reflect how much the results can be expected to generalize to other settings.
- It is fine to include aspirational goals as motivation as long as it is clear that these goals are not attained by the paper.

### 2. Limitations

Question: Does the paper discuss the limitations of the work performed by the authors? Answer: [Yes]

Justification: The paper includes a “Limitations and Future Work” section that discusses the limited availability of antigen-antibody paired data and the dependency on structural models for antigen encoding.

Guidelines:

- The answer NA means that the paper has no limitation while the answer No means that the paper has limitations, but those are not discussed in the paper.
- The authors are encouraged to create a separate “Limitations” section in their paper.
- The paper should point out any strong assumptions and how robust the results are to violations of these assumptions (e.g., independence assumptions, noiseless settings, model well-specification, asymptotic approximations only holding locally). The authors should reflect on how these assumptions might be violated in practice and what the implications would be.
- The authors should reflect on the scope of the claims made, e.g., if the approach was only tested on a few datasets or with a few runs. In general, empirical results often depend on implicit assumptions, which should be articulated.
- The authors should reflect on the factors that influence the performance of the approach. For example, a facial recognition algorithm may perform poorly when image resolution is low or images are taken in low lighting. Or a speech-to-text system might not be used reliably to provide closed captions for online lectures because it fails to handle technical jargon.
- The authors should discuss the computational efficiency of the proposed algorithms and how they scale with dataset size.
- If applicable, the authors should discuss possible limitations of their approach to address problems of privacy and fairness.
- While the authors might fear that complete honesty about limitations might be used by reviewers as grounds for rejection, a worse outcome might be that reviewers discover limitations that aren’t acknowledged in the paper. The authors should use their best judgment and recognize that individual actions in favor of transparency play an important role in developing norms that preserve the integrity of the community. Reviewers will be specifically instructed to not penalize honesty concerning limitations.

### 3. Theory Assumptions and Proofs

Question: For each theoretical result, does the paper provide the full set of assumptions and a complete (and correct) proof?

Answer: [NA]

Justification: The paper does not include theoretical results but focuses on the empirical evaluation of the proposed AIDA model for antibody design and optimization

Guidelines:

- The answer NA means that the paper does not include theoretical results.
- All the theorems, formulas, and proofs in the paper should be numbered and crossreferenced.
- All assumptions should be clearly stated or referenced in the statement of any theorems.
- The proofs can either appear in the main paper or the supplemental material, but if they appear in the supplemental material, the authors are encouraged to provide a short proof sketch to provide intuition.
- Inversely, any informal proof provided in the core of the paper should be complemented by formal proofs provided in appendix or supplemental material.
- Theorems and Lemmas that the proof relies upon should be properly referenced.

### 4. Experimental Result Reproducibility

Question: Does the paper fully disclose all the information needed to reproduce the main experimental results of the paper to the extent that it affects the main claims and/or conclusions of the paper (regardless of whether the code and data are provided or not)?

Answer: [Yes]

Justification: The paper provides comprehensive details on the datasets used, the experimental settings, and the evaluation metrics. It specifies the use of the SAbDab and SKEMPI V2.0 datasets, the pre-training process, and the exact steps taken for the antibody design tasks, ensuring reproducibility. Additionally, a GitHub repository is included for further access to the code and data.

Guidelines:

- The answer NA means that the paper does not include experiments.
- If the paper includes experiments, a No answer to this question will not be perceived well by the reviewers: Making the paper reproducible is important, regardless of whether the code and data are provided or not.
- If the contribution is a dataset and/or model, the authors should describe the steps taken to make their results reproducible or verifiable.
- Depending on the contribution, reproducibility can be accomplished in various ways. For example, if the contribution is a novel architecture, describing the architecture fully might suffice, or if the contribution is a specific model and empirical evaluation, it may be necessary to either make it possible for others to replicate the model with the same dataset, or provide access to the model. In general. releasing code and data is often one good way to accomplish this, but reproducibility can also be provided via detailed instructions for how to replicate the results, access to a hosted model (e.g., in the case of a large language model), releasing of a model checkpoint, or other means that are appropriate to the research performed.
- While NeurIPS does not require releasing code, the conference does require all submissions to provide some reasonable avenue for reproducibility, which may depend on the nature of the contribution. For example
  a. If the contribution is primarily a new algorithm, the paper should make it clear how to reproduce that algorithm.
  b. If the contribution is primarily a new model architecture, the paper should describe the architecture clearly and fully.
  c. If the contribution is a new model (e.g., a large language model), then there should either be a way to access this model for reproducing the results or a way to reproduce the model (e.g., with an open-source dataset or instructions for how to construct the dataset).
  d. We recognize that reproducibility may be tricky in some cases, in which case authors are welcome to describe the particular way they provide for reproducibility. In the case of closed-source models, it may be that access to the model is limited in some way (e.g., to registered users), but it should be possible for other researchers to have some path to reproducing or verifying the results.

### 5. Open access to data and code

Question: Does the paper provide open access to the data and code, with sufficient instructions to faithfully reproduce the main experimental results, as described in supplemental material?

Answer: [Yes]

Justification: The paper includes a GitHub repository link, providing access to the data and code used in the experiments, along with detailed instructions for reproducing the main experimental results.

Guidelines:

- The answer NA means that paper does not include experiments requiring code.
- Please see the NeurIPS code and data submission guidelines (https://nips.cc/public/guides/CodeSubmissionPolicy) for more details.
- While we encourage the release of code and data, we understand that this might not be possible, so “No” is an acceptable answer. Papers cannot be rejected simply for not including code, unless this is central to the contribution (e.g., for a new open-source benchmark).
- The instructions should contain the exact command and environment needed to run to reproduce the results. See the NeurIPS code and data submission guidelines (https://nips.cc/public/guides/CodeSubmissionPolicy) for more details.
- The authors should provide instructions on data access and preparation, including how to access the raw data, preprocessed data, intermediate data, and generated data, etc.
- The authors should provide scripts to reproduce all experimental results for the new proposed method and baselines. If only a subset of experiments are reproducible, they should state which ones are omitted from the script and why.
- At submission time, to preserve anonymity, the authors should release anonymized versions (if applicable).
- Providing as much information as possible in supplemental material (appended to the paper) is recommended, but including URLs to data and code is permitted.

### 6. Experimental Setting/Details

Question: Does the paper specify all the training and test details (e.g., data splits, hyperparameters, how they were chosen, type of optimizer, etc.) necessary to understand the results?

Answer: [Yes]

Justification: The paper specifies the datasets used (SAbDab and SKEMPI V2.0), the training and testing details, including data splits, hyperparameters, and the type of optimizer, necessary to understand the results.

Guidelines:

- The answer NA means that the paper does not include experiments.
- The experimental setting should be presented in the core of the paper to a level of detail that is necessary to appreciate the results and make sense of them.
- The full details can be provided either with the code, in appendix, or as supplemental material.

### 7. Experiment Statistical Significance

Question: Does the paper report error bars suitably and correctly defined or other appropriate information about the statistical significance of the experiments?

Answer: [Yes]

Justification: The paper reports error bars and statistical significance for the experiments, including the metrics used and the method for calculating these values.

Guidelines:

- The answer NA means that the paper does not include experiments.
- The authors should answer “Yes” if the results are accompanied by error bars, confidence intervals, or statistical significance tests, at least for the experiments that support the main claims of the paper.
- The factors of variability that the error bars are capturing should be clearly stated (for example, train/test split, initialization, random drawing of some parameter, or overall run with given experimental conditions).
- The method for calculating the error bars should be explained (closed form formula, call to a library function, bootstrap, etc.)
- The assumptions made should be given (e.g., Normally distributed errors).
- It should be clear whether the error bar is the standard deviation or the standard error of the mean.
- It is OK to report 1-sigma error bars, but one should state it. The authors should preferably report a 2-sigma error bar than state that they have a 96% CI, if the hypothesis of Normality of errors is not verified.
- For asymmetric distributions, the authors should be careful not to show in tables or figures symmetric error bars that would yield results that are out of range (e.g. negative error rates).
- If error bars are reported in tables or plots, The authors should explain in the text how they were calculated and reference the corresponding figures or tables in the text.

### 8. Experiments Compute Resources

Question: For each experiment, does the paper provide sufficient information on the computer resources (type of compute workers, memory, time of execution) needed to reproduce the experiments?

Answer: [Yes]

Justification: The paper provides detailed information on the compute resources used, including the type of GPU (NVIDIA A40), memory, and time of execution for each experiment.

Guidelines:

- The answer NA means that the paper does not include experiments.
- The paper should indicate the type of compute workers CPU or GPU, internal cluster, or cloud provider, including relevant memory and storage.
- The paper should provide the amount of compute required for each of the individual experimental runs as well as estimate the total compute.
- The paper should disclose whether the full research project required more compute than the experiments reported in the paper (e.g., preliminary or failed experiments that didn’t make it into the paper).

### 9. Code Of Ethics

Question: Does the research conducted in the paper conform, in every respect, with the NeurIPS Code of Ethics https://neurips.cc/public/EthicsGuidelines?

Answer: [Yes]

Justification: The research conforms to the NeurIPS Code of Ethics, with no ethical concerns identified in the paper.

Guidelines:

- The answer NA means that the authors have not reviewed the NeurIPS Code of Ethics.
- If the authors answer No, they should explain the special circumstances that require a deviation from the Code of Ethics.
- The authors should make sure to preserve anonymity (e.g., if there is a special consideration due to laws or regulations in their jurisdiction).

### 10. Broader Impacts

Question: Does the paper discuss both potential positive societal impacts and negative societal impacts of the work performed?

Answer: [Yes]

Justification: The paper discusses the potential positive impacts of improving antibody design and optimization, as well as addressing potential negative impacts, such as the misuse of the technology for malicious purposes.

Guidelines:

- The answer NA means that there is no societal impact of the work performed.
- If the authors answer NA or No, they should explain why their work has no societal impact or why the paper does not address societal impact.
- Examples of negative societal impacts include potential malicious or unintended uses (e.g., disinformation, generating fake profiles, surveillance), fairness considerations (e.g., deployment of technologies that could make decisions that unfairly impact specific groups), privacy considerations, and security considerations.
- The conference expects that many papers will be foundational research and not tied to particular applications, let alone deployments. However, if there is a direct path to any negative applications, the authors should point it out. For example, it is legitimate to point out that an improvement in the quality of generative models could be used to generate deepfakes for disinformation. On the other hand, it is not needed to point out that a generic algorithm for optimizing neural networks could enable people to train models that generate Deepfakes faster.
- The authors should consider possible harms that could arise when the technology is being used as intended and functioning correctly, harms that could arise when the technology is being used as intended but gives incorrect results, and harms following from (intentional or unintentional) misuse of the technology.
- If there are negative societal impacts, the authors could also discuss possible mitigation strategies (e.g., gated release of models, providing defenses in addition to attacks, mechanisms for monitoring misuse, mechanisms to monitor how a system learns from feedback over time, improving the efficiency and accessibility of ML).

### 11. Safeguards

Question: Does the paper describe safeguards that have been put in place for responsible release of data or models that have a high risk for misuse (e.g., pretrained language models, image generators, or scraped datasets)?

Answer: [NA]

Justification: The paper does not pose high risks for misuse, thus no specific safeguards are necessary.

Guidelines:

- The answer NA means that the paper poses no such risks.
- Released models that have a high risk for misuse or dual-use should be released with necessary safeguards to allow for controlled use of the model, for example by requiring that users adhere to usage guidelines or restrictions to access the model or implementing safety filters.
- Datasets that have been scraped from the Internet could pose safety risks. The authors should describe how they avoided releasing unsafe images.
- We recognize that providing effective safeguards is challenging, and many papers do not require this, but we encourage authors to take this into account and make a best faith effort.

### 12. Licenses for existing assets

Question: Are the creators or original owners of assets (e.g., code, data, models), used in the paper, properly credited and are the license and terms of use explicitly mentioned and properly respected?

Answer: [Yes]

Justification: The paper properly credits the creators of the datasets and models used, and mentions the licenses and terms of use explicitly.

Guidelines:

- The answer NA means that the paper does not use existing assets.
- The authors should cite the original paper that produced the code package or dataset.
- The authors should state which version of the asset is used and, if possible, include a URL.
- The name of the license (e.g., CC-BY 4.0) should be included for each asset.
- For scraped data from a particular source (e.g., website), the copyright and terms of service of that source should be provided.
- If assets are released, the license, copyright information, and terms of use in the package should be provided. For popular datasets, paperswithcode.com/datasets has curated licenses for some datasets. Their licensing guide can help determine the license of a dataset.
- For existing datasets that are re-packaged, both the original license and the license of the derived asset (if it has changed) should be provided.
- If this information is not available online, the authors are encouraged to reach out to the asset’s creators.

### 13. New Assets

Question: Are new assets introduced in the paper well documented and is the documentation provided alongside the assets?

Answer: [Yes]

Justification: The new assets introduced in the paper are well documented, and the documentation is provided alongside the assets in the supplemental material and GitHub repository.

Guidelines:

- The answer NA means that the paper does not release new assets.
- Researchers should communicate the details of the dataset/code/model as part of their submissions via structured templates. This includes details about training, license, limitations, etc.
- The paper should discuss whether and how consent was obtained from people whose asset is used.
- At submission time, remember to anonymize your assets (if applicable). You can either create an anonymized URL or include an anonymized zip file.

### 14. Crowdsourcing and Research with Human Subjects

Question: For crowdsourcing experiments and research with human subjects, does the paper include the full text of instructions given to participants and screenshots, if applicable, as well as details about compensation (if any)?

Answer: [NA]

Justification: The paper does not involve crowdsourcing or research with human subjects. Guidelines:

- The answer NA means that the paper does not involve crowdsourcing nor research with human subjects.
- Including this information in the supplemental material is fine, but if the main contribution of the paper involves human subjects, then as much detail as possible should be included in the main paper.
- According to the NeurIPS Code of Ethics, workers involved in data collection, curation, or other labor should be paid at least the minimum wage in the country of the data collector.

### 15. Institutional Review Board (IRB) Approvals or Equivalent for Research with Human Subjects

Question: Does the paper describe potential risks incurred by study participants, whether such risks were disclosed to the subjects, and whether Institutional Review Board (IRB) approvals (or an equivalent approval/review based on the requirements of your country or institution) were obtained?

Answer: [NA]

Justification: The paper does not involve crowdsourcing or research with human subjects. Guidelines:

- The answer NA means that the paper does not involve crowdsourcing nor research with human subjects.
- Depending on the country in which research is conducted, IRB approval (or equivalent) may be required for any human subjects research. If you obtained IRB approval, you should clearly state this in the paper.
- We recognize that the procedures for this may vary significantly between institutions and locations, and we expect authors to adhere to the NeurIPS Code of Ethics and the guidelines for their institution.
- For initial submissions, do not include any information that would break anonymity (if applicable), such as the institution conducting the review.

## Notes

### Competing Interest Statement

The authors have declared no competing interest.

### Summary of Updates

In this revised version of the manuscript, we addressed inconsistencies between data and explanations in the ablation study section. We ensured that the results presented in the tables align accurately with the corresponding analysis providing a clearer and more accurate representation of the impact of each model component on AIDA's performance across different antibody regions. This revision enhances the manuscript's coherence and reliability, making it easier for readers to follow the logical progression of our findings. By directly addressing these discrepancies, we reinforce the validity of our results and improve the clarity of the manuscript overall.

